# Resolving cellular systems by ultra-sensitive and economical single-cell transcriptome filtering

**DOI:** 10.1101/800631

**Authors:** Andres F. Vallejo, James Davies, Amit Grover, Ching-Hsuan Tsai, Robert Jepras, Marta E. Polak, Jonathan West

**Author notes:** Joint senior author (Marta E. Polak, Jonathan West). Correspondence to: Dr Jonathan West, Dr Marta E Polak. **Authors contributions:** AFV, MEP and JW conceived the idea, planned the experiments and wrote the manuscript. AFV, JD and AG performed the experiments. AFV and MEP analysed the data. AG, CHT and RJ contributed to the experimental plan and reviewed the manuscript.

## Abstract

Single-cell transcriptomics has sensitivity limits that restrict low abundance transcript identification, affects clustering and introduce artefact. Here, we describe Constellation DropSeq (C-DropSeq), a molecular transcriptome filter that delivers two orders of magnitude sensitivity gains by maximising read utility while reducing sequencing depth and costs. The simple and powerful method is broadly compatible with library preparation routines and was demonstrated by identifying and characterizing the activation of rare dendritic cell sub-populations.

## Main

The dramatic uptake and expansion of single-cell transcriptome analysis tools has transformed biological research, enabling reconstruction of population architectures and underlying processes to be revealed. The tools rely on compartmentalisation of single cells with the introduction of unique genetic barcodes during library preparation^1^. Though formidable, not unexpectedly these methods have sensitivity limits, with associated transcript absence events (dropouts) that restrict the faithful delineation of cell subtypes and especially overlook low abundant transcripts such as transcription factors, receptors and signalling molecules that are often pivotal for accurately describing cell processes and fate^2,3^. This is a consequence of high abundance transcripts occupying the available NGS read space and exacerbated by exponential PCR-directed library preparation routines.

Targeted approaches forgo global transcriptome screens, preferring to select transcripts of known utility and are especially favoured for mechanistic studies. Diverse targeted strategies have emerged; physical recovery of transcriptome subsets^4^, coupling custom primers to poly(dT) capture beads (DART-seq)^5^ and panel selection by PCR as with the Rhapsody workflow (BD)^6^. These methods are technically challenging and introduce substantial costs. Here, we describe Constellation DropSeq (C-DropSeq), a remarkably simple, inexpensive and scalable (e.g. >200 targets) approach, introducing a linear amplification stage in advance of conventional library preparation. Superior performance is demonstrated with two orders of magnitude sensitivity gains using only a 1/10^th^ of the sequencing depth for describing system architectures and processes with unprecedented resolution.

The DropSeq beads support 10^10^ probes^5^ indicating that sensitivity losses arise from the restricted NGS read space (~10^4–6^/cell) and also from exponential PCR amplification during library preparation, where abundant and more efficiently replicated transcripts dominate the available reads. In contrast, linear (single primer) amplification provides an unbiased route to enrichment across transcripts^7,8^. Therefore, in our C-DropSeq approach we have used linear amplification following cDNA synthesis for the targeted enrichment of transcripts of interest. The method involves replacing the template switching oligo (TSO) with hybrid primers containing a transcript-specific region adjacent to a universal handle to select and barcode desired transcripts in a single linear amplification (Fig. 1A; supplementary Fig. 1). We introduced this linear targeted amplification step to the DropSeq pipeline to provide a direct comparison that is amenable to cost-effective, large-scale cell screening campaigns albeit with recognised dropout limitations^1^.

**Figure 1.**
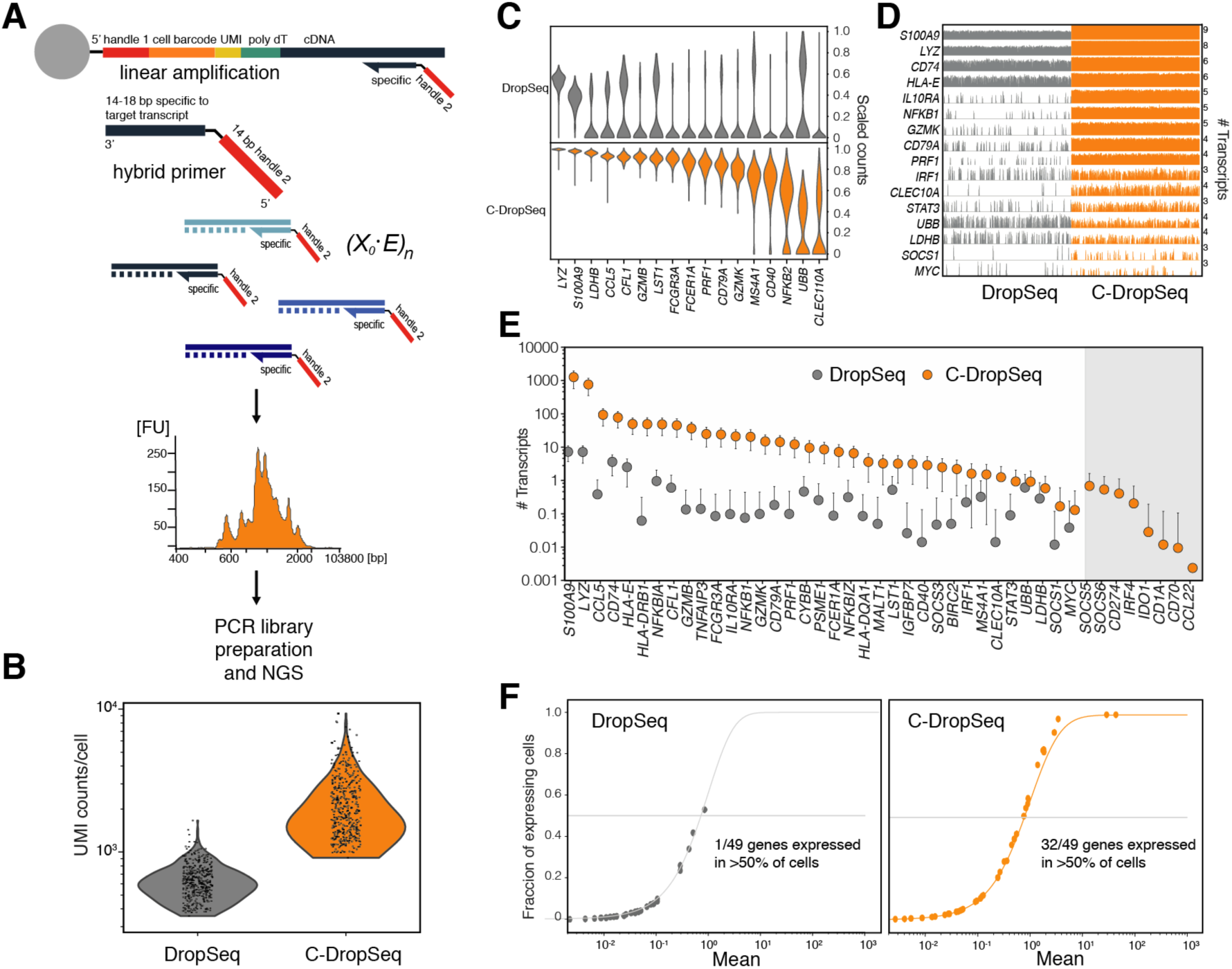
Characterisation of Constellation DropSeq. A) C-DropSeq protocol: A hybrid primer (14-18 bp specific sequence, black, adjacent to a common 14 bp handle 2, red) binds to a specific target sequence of cDNA captured on Macosko beads^9^. A panel of such hybrid primers can be introduced to standard DropSeq pipeline following cell encapsulation and generation of STAMPS. Linear amplification of 500-1000 bp stretches of target transcripts allows selective enrichment of targets of interest, and the inclusion of the cell barcode and UMI sequences, leads to generation of constellation library, ready to use in Next Generation Sequencing. The initial template copy number *X*_*o*_ multiplied by the replication efficiency *E* and the cycle number *n*. B-F) C- DropSeq was compared against DropSeq using a panel of 52 targets with control beads. B) C-DropSeq UMI counts per bead are 2.7-fold greater than with DropSeq. C) Scaled individual target transcripts counts show ~100-fold sensitivity gains for genes selected in C-DropSeq. D) A trackplot showing the data structure in a head to head comparison. Each bar represents a gene expression signal from a single cell. Full Trackplot is included as Supplementary Fig. 4. E) The fraction of expressing beads as a function of the mean expression was used as the comparator, error bars represent SD. F) Dramatic reduction in dropouts achieved by C-DropSeq compared with DropSeq. At 2K UMI counts per bead 32/49 genes were detected in half of the beads (3 negative controls were not detected) in C-DropSeq whereas only 1 was detected with the same threshold in DropSeq.

Assay development first involved a panel of 20 primers applied to control beads^9,10^ bearing a bulk RNA sample to exclude biological variation. Sensitivity was compared between single primer linear amplification and dual primer exponential amplification (PCR, requiring a SMART-Seq reverse primer) akin to state of the art methods (*e.g.* Rhapsody, BD)^6^. The primer panel contained high, medium and low expression level transcripts specific for peripheral blood mononuclear cells (PBMCs) and activation traits (**Supplementary Table 1**). C-DropSeq is amplification cycle and primer concentration dependent method (Supplementary Fig. 2), with straightforward optimisation enabling the selective capture of desired transcripts which produce a characteristically spiny tapestation plot (Fig. 1A). Critically, at 12K reads/bead, linear amplification has a low, 6.8 duplication rate, producing 1,818 UMIs per bead to enable the detection of 17/20 transcripts using a 50% dropout cut-off. In contrast, exponential amplification has a 24.1 duplication rate, reducing the UMI number to 467 and resulting in only 13/20 transcripts attaining the 50% dropout cut-off (Supplementary Fig. 3).

Next, C-DropSeq was scaled to 52 targets including 3 negative controls and compared with standard DropSeq(**Supplementary Table 2**). Using 15k reads/bead, we demonstrated efficient use of the read space (93.5% reads) and increasing the average UMI counts/cell at 2.7-fold (Fig. 1B). Individual target transcript counts from C-DropSeq were on average 83-fold higher (Fig. 1C), dramatically reducing the dropout rate and providing a uniform gene expression distribution to accurately rank expressed transcripts (Fig.1 C,D; Supplementary Fig. 5). Standard DropSeq only detected 41 of the targets, while C-DropSeq detected all 49 targets and none of the control genes (Fig. 1E). The 8 transcripts exclusively detected by C-DropSeq had average expression levels ranging from 0.03–2.60 counts per ten thousand (CPTT), without length correlation. In practical terms, when using a 50% dropout cut-off, 32/49 are detected by C-DropSeq and only 1/49 by standard DropSeq at a sequencing depth of 8k reads/bead (Fig. 1F). Of merit, the sensitivity of C-DropSeq cascades directly into significantly lower read requirements; the 32/49 transcripts above 50% cut-off are detected when reducing the depth to 4k reads/bead, with losses (28/49) only evident at 2k (Supplementary Fig. 5). This striking feature of C-DropSeq presents the option to reduce the sequencing depth and associated experimental cost, or increase the scale of the experiment.

To explore the ability of C-DropSeq to measure gene expression changes in response to perturbation in a cellular system, we challenged human peripheral blood mononuclear cells (PBMCs) with a super antigen Staphylococcal enterotoxin B (SEB, 100 ng/mL, 16 hours). To compare methods 1000 cells per treatment were sequenced (DropSeq: 200K reads/cells C-DropSeq: 20Kreads/cell, Fig. 2A). In this context, C-DropSeq consistently detected low copy transcripts such as *GZMB, IRF4* and *SOCS1* with reduced dropout and increased UMI counts at 10-fold lower sequencing depth. Differential gene expression was compared between control and stimuli for both standard DropSeq and C-DropSeq. The fold change measurements correlated well between the methods (r=0.62, *p*-value=8e-5, Fig. 2B,C). Importantly, C-DropSeq was 1.6 times more sensitive (assessed by gradient) to gene expression changes (Fig. 2B), improving the resolution of typical activation features such as *NFKB1/NFKBIA* while maintaining comparable expression levels for stable transcripts unperturbed by stimulation (*e.g. CD74*). In summary, the linear amplification step in C-DropSeq retains the authentic biological response, while measuring responses with greater sensitivity and resolving greater detail in the underlying process (Supplementary Fig. 5).

**Figure 2.**
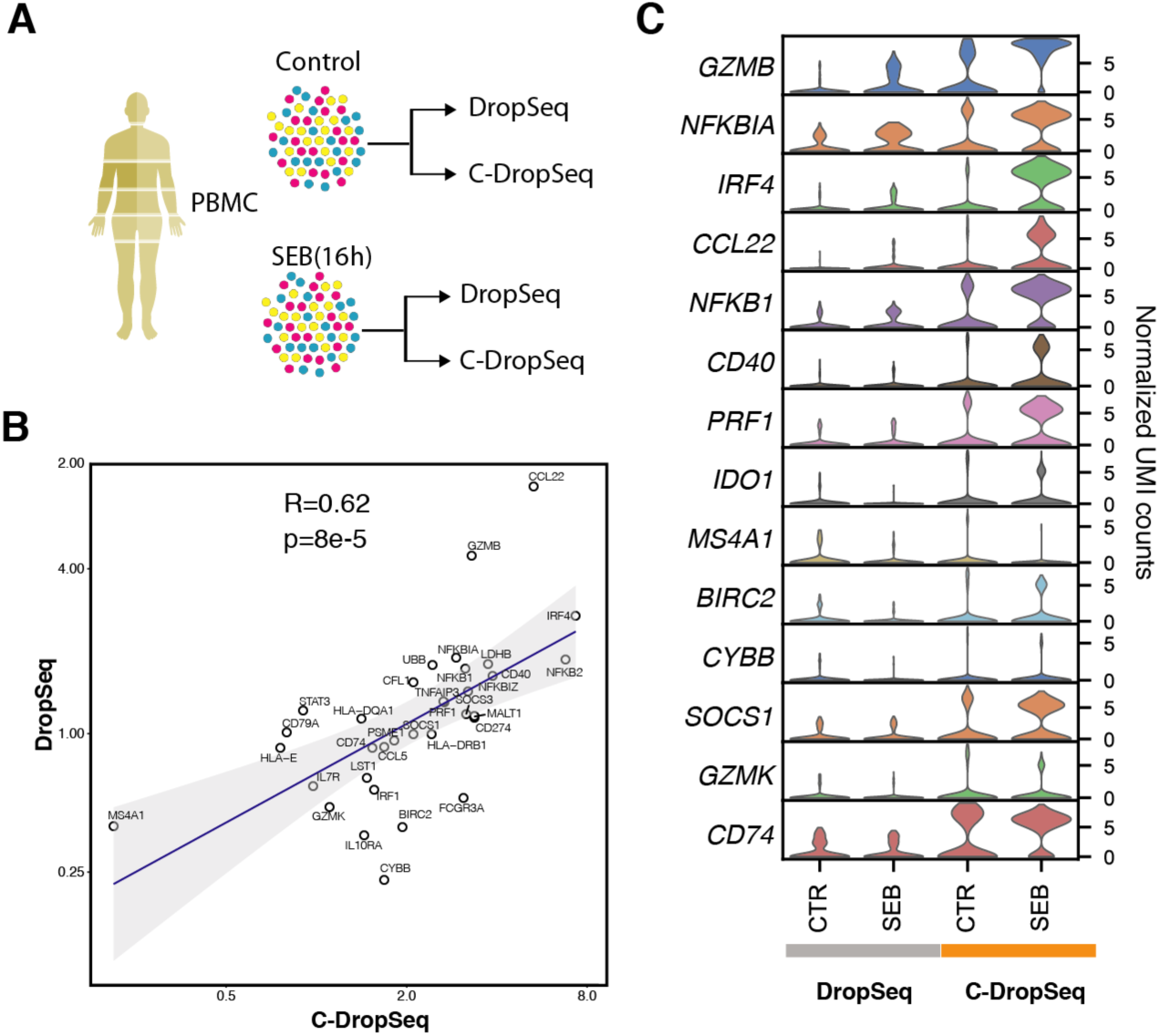
Constellation DropSeq reliably measures changes in gene expression. A) Experimental design; PBMCs from healthy subjects (n=3) were stimulated with Staphylococcal enterotoxin B (SEB) or media control for 16h and analysed using DropSeq and C-DropSeq. B) Correlation of normalised gene expression fold changes induced by SEB as detected by DropSeq and C-DropSeq. Pseudo-bulk counts for each gene used for the comparison. C) Comparative analysis of selected markers induced by SEB in cultured PBMCs. Violin plots in each row show the distribution and levels of each expressed gene in different culture conditions (CTR – media control, SEB – stimulated cells) and assessed by DropSeq (grey) and C-DropSeq (orange). y axis represents normalized UMI counts.

To demonstrate the applicability of C-DropSeq for the analysis of specific cell subtypes within complex cellular systems, we designed a primer panel targeting 127 transcripts (**Supplementary Table 3**) using a recent molecular classification^11^ for the identification of dendritic cell (DC) subpopulations and their activation states. 4000 Human PBMCs were cultured with Gram-negative bacterial endotoxin lipopolysaccharide (LPS, 1 μg/mL for 4 hours), a potent inflammatory mediator inducing DC activation via TLR4. While standard DropSeq was able to segregate the blood cell types, including DCs and monocytes (Fig. 3A), the technique was not sufficiently sensitive to detect DC sub-populations. In contrast, the sensitivity of C-DropSeq allowed the classification of expression markers for two DC subpopulations (DC1: *CLEC9A*, and DC2: *FCERIA*^11^ Supplementary Fig.6, Fig. 3B), Furthermore, C-DropSeq provided greater insights into LPS-induced transcriptome remodelling, including the identification of activation markers (*CCR7*), cytokines (*IL1B*) and low abundance transcription factors (*IRF7*,) in the DC1 subpopulation (Fig. 3C), and up-regulation of *CD83*, *CCR7* and *PSMA1* in DC2 (Supplementary Fig. 6).

**Figure 3.**
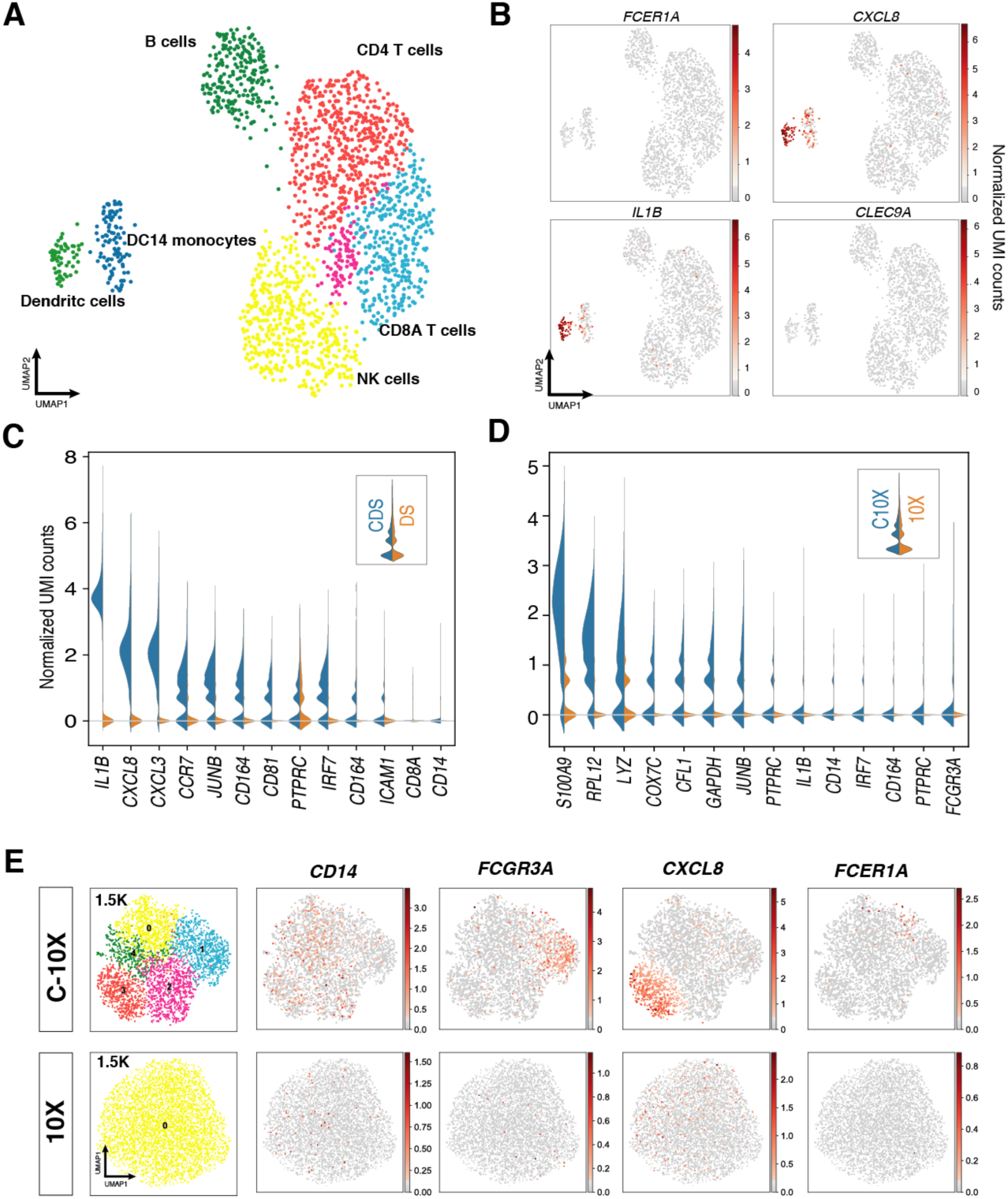
Constellation DropSeq can resolve rare cell populations. DropSeq assay of human PBMC stimulated with lipopolysaccharide (LPS). **A)** Control and 4 hours LPS stimulated PBMC were analysed using DropSeq. UMAP plot of the merged dataset is shown (2542 single cells). Expression of selected markers induced in response to LPS in DCs and monocytes and DC-subset specific markers (red) (Scanpy, UMAP plot: Leiden r = 0.5, n_pcs=10, n_neighbours =20). **B)** Expression of low abundance genes in DC population by DropSeq. Markers for DC1 (*CLEC9A*) and DC2 (*FCER1A*)^11^ and are not detected. **C)** Direct comparison of differential gene expression in dendritic cell population in response to LPS Stimuli using C-DropSeq vs DropSeq. Left: C-DropSeq (blue) vs DropSeq (orange), gene expression measured in normalised UMAP counts D) Direct comparison of gene expression in monocytes using C-10X vs 10X. Left: C-10X (blue) vs 10X (orange), gene expression measured in normalised UMAP counts E). UMAP plot of 6000 monocyte transcriptomes assessed using 10X and C-10X. At 1.5K UMI counts per cell, C-10X shows more granularity than normal 10X at the same resolution. The enhanced sensitivity of C-10X is represented using monocyte markers.

Next the constellation approach was reconfigured for use with the popular Chromium 10x Genomics technology using 6000 CD14 enriched monocytes. Constellation-10X (C-10X) greatly improved the detection of transcripts of interest (Fig.3D). C-10X showed 22-fold greater sensitivity allowing reduction of the sequencing depth from 70k to 1.5K reads/cell, while distinguishing 5 clusters and identifying rare dendritic cells (*FCER1A*) and an activated monocyte sub-population (*CXCL8*). In comparison standard 10X at 1.5K reads/cell failed to resolve these sub-populations and activation states (Fig. 3E, Supplementary Fig.7). Indeed, standard 10X requires 70k reads/cell to obtain the same results, inflating the experimental costs 46-fold (Supplementary Fig. 7) demonstrating both the sensitivity and financial gains achieved using the Constellation approach.

The simplicity of the Constellation method allows inclusion in almost any single cell transcriptome library preparation pipelines involving SMART-Seq primers (DropSeq, Seq-Well, 10x and potentially InDrop). The multiplex scaling capacity is governed by available volume; a 300-plex assay is feasible for a 50 μL reaction volume (without affection the normal library preparation pipeline; Supplementary Fig. 8). The highly multiplexed selection of transcripts of interest is at the expense of global transcriptome coverage, yet benefits from maximising the efficient use of the NGS space to enable ultra-sensitive investigations. In this manner, the architecture of cellular systems can be understood with unprecedented resolution and biological processes can be mapped in exquisite detail. Central to C-DropSeq is prior knowledge of the cellular system, where specific target selection lends strength to mechanistic studies or allows the prioritisation of targets for targeted perturbation. Additionally, C-DropSeq can be implemented efficiently in drug discovery, and preliminary toxicity and efficacy screens for pharmacological compounds of interest. To gain entry to new biological scenarios and to define the targeted primer library for C-DropSeq, various standard scRNA-seq approaches or bulk transcriptome analyses can first be applied to provide a global screen of the defining molecules and pathways of interest.

C-DropSeq builds on standard DropSeq, an already cost-effective single cell transcriptomics approach for large-scale experiments, while addressing the issues of sensitivity and dropout. With C-DropSeq further savings emerge from shrinking the required sequencing depth to allow substantially larger experiments or simply more experiments. The experimental economies, including time-finance trade-offs, of C-DropSeq and C-10x are compared with standard DropSeq and 10x approaches in **Supplementary Table 4** to inform method selection by end-users. Beyond this, C-DropSeq is accessible to resource limited laboratories, overall representing a step towards the democratisation of single-cell transcriptomics and the broad-scale expansion of our understanding of biological systems.

## Methods

### Primer Design

Primers targeting genes of interest were designed using Beacon Designer primer design software (PREMIER Biosoft, California US). The last 14 bases from the SMART primer sequence (TATCAACGCAGAGT) were added to the 5’ end of the designed primers. Desired features of primers included: a length between 28-32 base pairs, 40-60% GC content, a primer melting temperature between 52-58°C, and with minimal chance of secondary structures being produced.

### Negative control beads

RNA from fresh PBMC was extracted using RNeasy Plus Mini Kit (Qiagen). Control beads were generated by adding a solution of PBMC RNA at 10 pg/bead, making the RNA content in each droplet equivalent. 200 μL of reverse transcriptase mix (75 μL water, 40 μL Maxima 5x RT buffer, 40 μL 20% Ficoll PM-400, 20 μL 10 mM dNTPs, 5 μL RNase inhibitor and 10 μL Maxima H-RTase) was added to each bead sample. 10 μL of 50 μM TSO was added to the DropSeq controls, whereas for C-DropSeq no TSO was used. Samples were incubated with rotation at room temperature for 30 minutes followed by 90 minutes at 42°C with continuous rotation. Beads were washed with 1 mL TE-SDS (10 mM Tris, pH 8.0, 1 mM EDTA, 5% SDS) and twice with 1 mL TE-TW (10 mM Tris, pH 8.0, 1 mM EDTA, 0.01% Tween-20). Finally, beads were washed with 1 mL 10 mM Tris pH 8.0, and stored at 4C.

### Cell preparation

Human blood was collected from donors with written consent and ethical approval (study number: 17/EM/0349). PBMC were extracted immediately using Lymphoprep™ (STEMCELL Technologies) and incubated at 37°C with 5% CO_2_. For SEB stimulation experiments cells were cultured in 24 well plates at 2×10^6^ cells/mL for 16h with or without SEB, using a final SEB concentration of 100 ng/mL. For LPS stimulation experiments cells were cultured in 24 well plates at 2×10^6^ cells/mL for 4h with or without LPS, using a final LPS concentration of 1 μg/mL. Following the incubation period cells were harvested, washed in PBS and counted. 180,000 cells were taken for encapsulation. CD14+ monocytes for the 10X experiment were purchased from Tissue solutions (Glasgow, UK).

### DropSeq

Drop-Seq library preparation and sequencing was performed as described previously^9^. Briefly, single cells were co-encapsulated with beads in droplets using the microfluidic design provided by Macosko *et al*^9^. After cell lysis, cDNA synthesis was carried out (Maxima Reverse Transcriptase, Thermo Fisher), followed by PCR (Kapa Hotstart Ready mix, 15 cycles: 4 at 67C, 11 at 65C). cDNA libraries were tagmented and PCR-amplified (Nextera tagmentation kit, Illumina). Finally, libraries were pooled and sequenced on an Illumina Nextseq500, (paired end 20×50bp reads).

### Constellation DropSeq

For Constellation DropSeq, experiments were processed as normal from encapsulation through to extraction and purification of beads from the droplet emulsion. During reverse transcription however, the template switching oligo (TSO) was absent from the reaction*. This resulted in cDNA fragments without SMART primer binding sites at the 3’ end of the Macosko bead primers,. Hybrid primers were pooled at 10 μM. A 50 μL amplification mix was added (25 μL 2X Kapa HiFi Hotstart Readymix, 10 μM primer pool, 24.6 μL water) to aliquots of 2000 beads (~100 STAMPs). 20 rounds of linear amplification (at 60°C) were first performed before continuing the standard Drop-Seq protocol for library preparation with PCR amplification and tagmentation. cDNA libraries were purified twice using AMPure XP magnetic beads (Beckman Coulter) (1:0.6) and libraries assessed using Agillent bioanalyser (KIT) before tagmentation and Next-seq sequencing.

*Standard reagents including the TSO can be used with the caveat of transcript noise generated by the reverse SMART primer.

### 10X Genomics

Single cell libraries were generated using the Chromium Single Cell 3ʹ library and gel bead kit v3.1 from 10x Genomics. Briefly, 10,000 cells were loaded onto a channel of the 10x chip to produce Gel Bead-in-Emulsions (GEMs). This underwent reverse transcription to barcode RNA before clean-up and cDNA amplification followed by enzymatic fragmentation and 5ʹ adaptor and sample index attachment using the Nextera XT Library preparation kit (Illuumina). Libraries were sequenced on the MiSeq500 (Illumina) with 28×60 bp paired-end sequencing.

### Constellation 10X

For Constellation 10X, 395 pg of cDNA were used for linear amplification comprising 20 rounds of linear amplification (60°C) using a pool of primers at 10 μM. A 40 μL amplification mix was added (20 μL 2X Kapa HiFi Hotstart Readymix, 10 μM primer pool) to 10 μL of cDNA library. cDNA libraries were purified twice using AMPure XP (Beckman Coulter) magnetic beads (1:0.6) and libraries assessed using a bioanalyser before tagmentation and Next-seq sequencing on an Illumina Nextseq500, (paired end 28×60bp reads).

### Real Time PCR

Control beads were used to assess the specificity of C-DropSeq. 400 control beads per well were used as starting material. C-DropSeq libraries were produced by linear amplification using two control primers (CFL1 and UBB from IDT) for 5, 10 or 20 cycles. Libraries were purified twice using 0.6X AMPure XP magnetic beads (Beckman Coulter) and eluted with 20 μL 1xTE, pH 8.0. C-DropSeq libraries were tested using specific primers designed within the amplicon region including a negative control, CD74. 2 μL of the C-DropSeq library was amplified in iTaq™ Universal SYBR (Bio-Rad) containing 200 nM of CFL1, UBB or CD74 primers. Amplification was undertaken in technical triplicates on a HT7900 Fast Real-Time PCR System (Applied Biosystems). Quantification was achieved against a serial dilution calibration curve of the pool of samples in each plate. C_t_ values were thresholded at 0.1 relative fluorescence units (RFU).

### Bioinformatic pipelines

Alignment, read filtering, barcode and UMI counting were performed using kallisto-bustools^12^. High quality barcodes were selected based on the overall UMI distribution using emptyDrops^13^. All further analyses were run using the Python-based Scanpy^14^. To remove low quality cells, we filtered cells with a high fraction of counts from mitochondrial genes (20% or more) indicating stressed or dying cells^9^. In addition, genes expressed in less than 20 cells were excluded.

Cell by gene count matrices of all samples were concatenated to a single matrix and values log transformed. To account for differences in sequencing depth or cell size UMI counts were normalized using quantile normalization. The top variable genes were selected based on normalized dispersion. This output matrix was input to all further analyses except for differential expression testing where all genes were used.

### Visualization and clustering

A single-cell neighbourhood graph was computed on the 50 first principal components that sufficiently explain the variation in the data using 20 nearest neighbours. Uniform Manifold Approximation and Projection (UMAP) was run for visualization. For clustering and cell type identification Leiden-based clustering ^15^ at 0.5 resolution was used. Cell types were annotated based on the expression of known marker genes.

## Acknowledgements

We are grateful to the subjects who participated in this study; and Elena Vataga, Computational Modelling Group, University of Southampton, for assistance with the High-Performance Computing | HPC Platform.

## Ethics declarations

The PBMCs used in this study were obtained with ethical approval 17/EM/0349.

---

**Figure S1.**
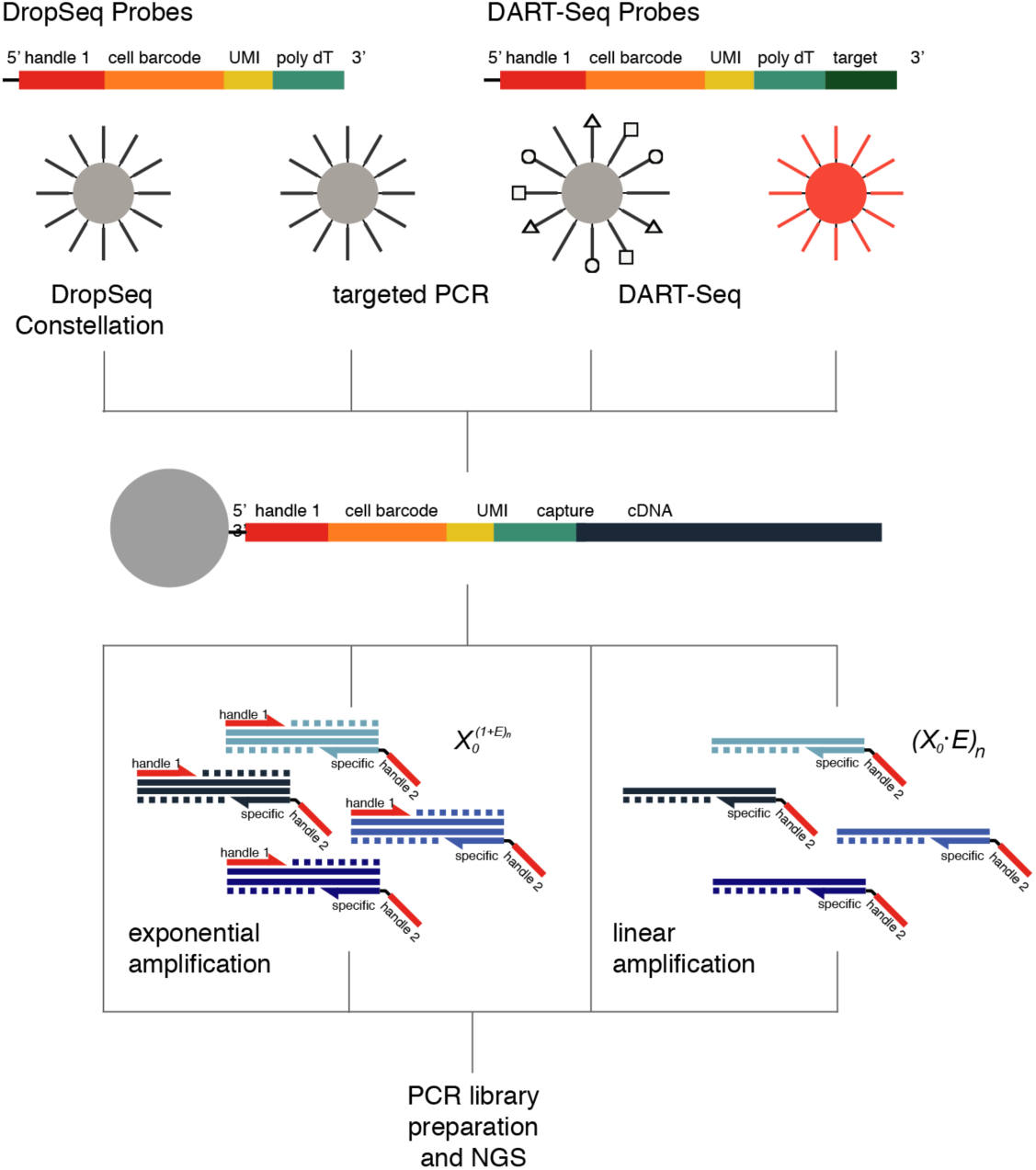
Comparison of the standard DropSeq with C-DropSeq and other targeted methods. The methods use the same capture probes, with the exception of the DART-Seq method that have probes extended with target-specific capture sequences (multiplexing is illustrated with shapes). Following cDNA capture DropSeq and DART-Seq methods progress directly to PCR library preparation, whereas targeted PCR and the Constellation Drop-Seq methods first involve PCR and linear amplification cycles, respectively. The amplification behaviour, exponential or linear, is described by the initial target number *X*_*o*_, the efficiency of replication *E* and the cycle number *n*.

**Figure S2.**
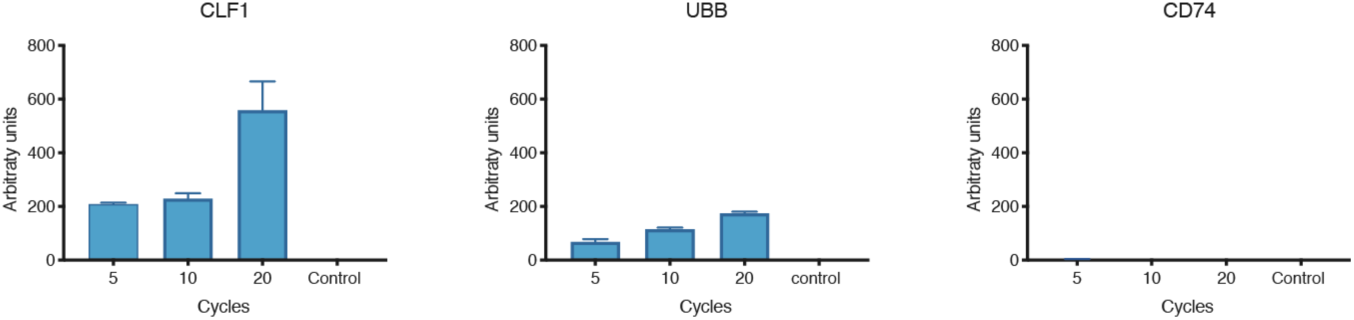
Expression analysis of a C-DropSeq library containing CLF1 and UBB primers. The library generated from control beads using linear amplification, at primer concentration10 nMol and 65°C annealing temperature was tested with qPCR for expression of *CLF1* and *UBB* as targeted genes and *CD74* as a negative control. Error bars represent standard deviation (SD).

**Figure S3.**
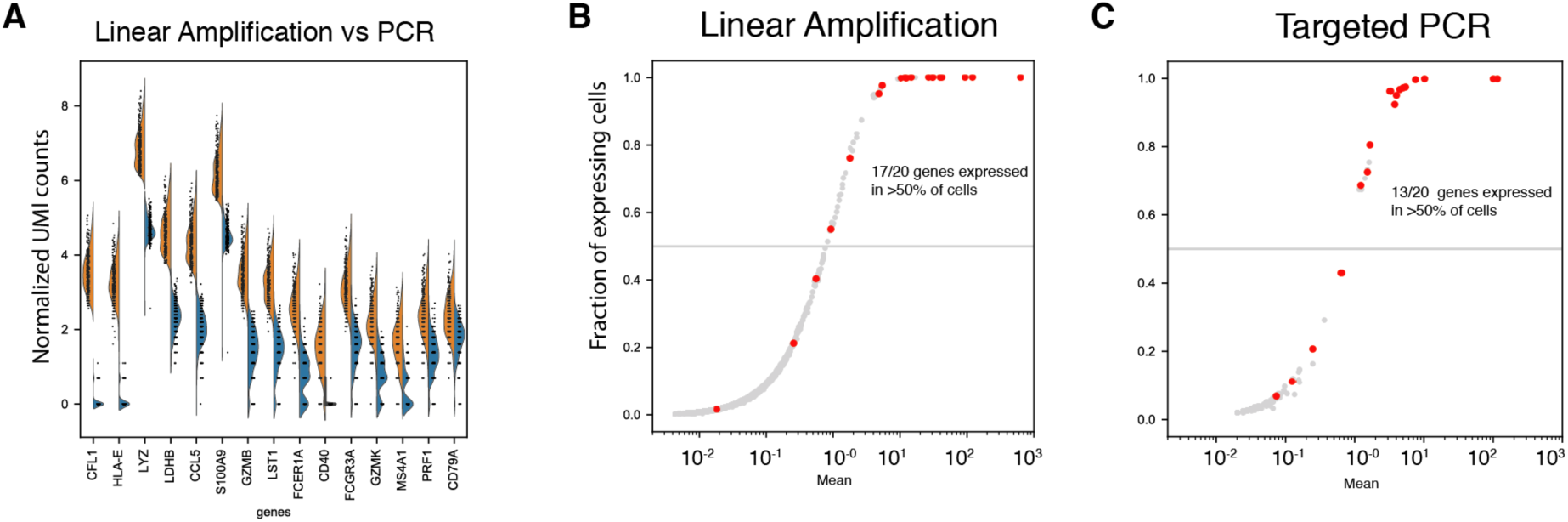
Head to head comparison of detection of 20 targets using linear vs targeted approach. A) Expression levels (Normalized UMI counts) detected by linear amplification (orange) and PCR (blue). B) Drop-out rate vs mean expression levels in linear amplification. Red dots represent genes included in test library. C) Drop-out rate vs mean expression levels in targeted PCR. Red dots represent genes included in the library tested.

**Figure S4.**
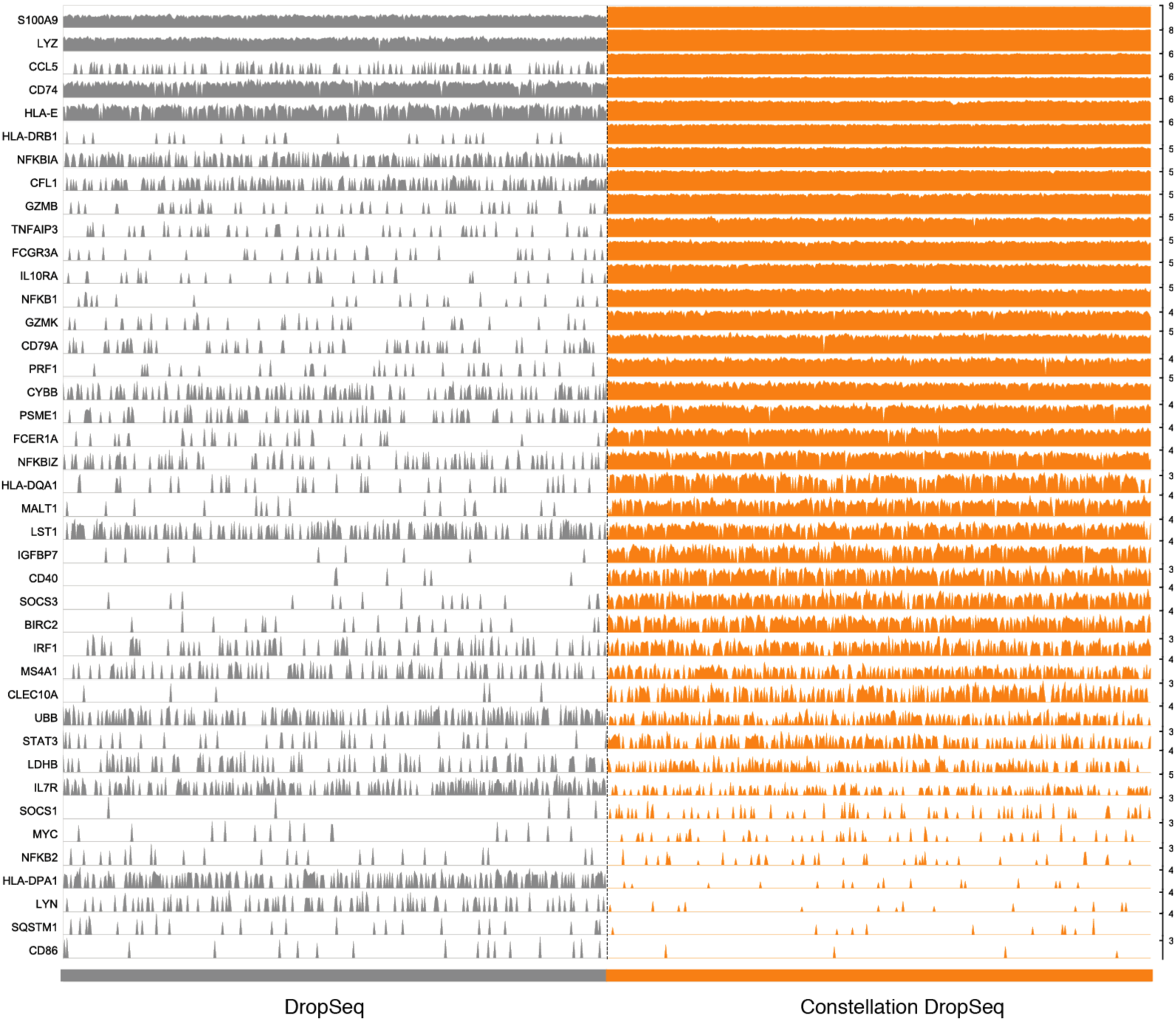
DropSeq and C-DropSeq comparison for the detection of a panel of 52 genes. Trackplot of gene expression for high, medium and low expressed genes detected using Drop-seq (grey) and C-DropSeq (orange) with control beads. A total of 41/52 genes were detected in both methods. Each bar shows the UMI counts signal from a single cell.

**Figure S5.**
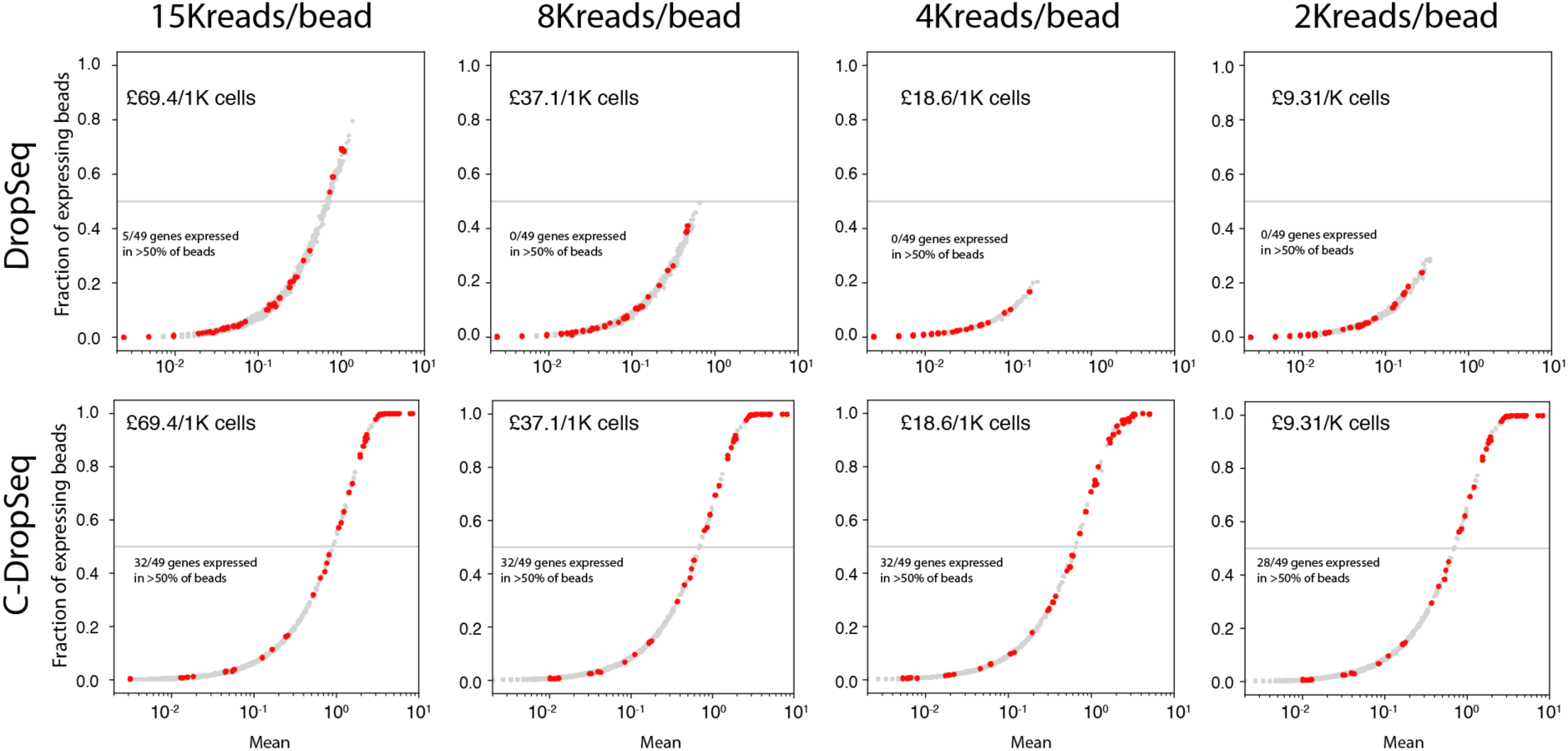
DropSeq and C-DropSeq sensitivity comparison varying sequencing depth. The total number of counts for each target was calculated and compared between DropSeq (top) and C-DropSeq (bottom). The fraction of beads with detected target expression vs mean level of target expression are shown for each gene. The horizontal line indicates the 50% of beads detection threshold Red: genes from the panel. Grey: genes not included in the panel. Numbers are the predicted effective cost for 1000 cells.

**Figure S6.**
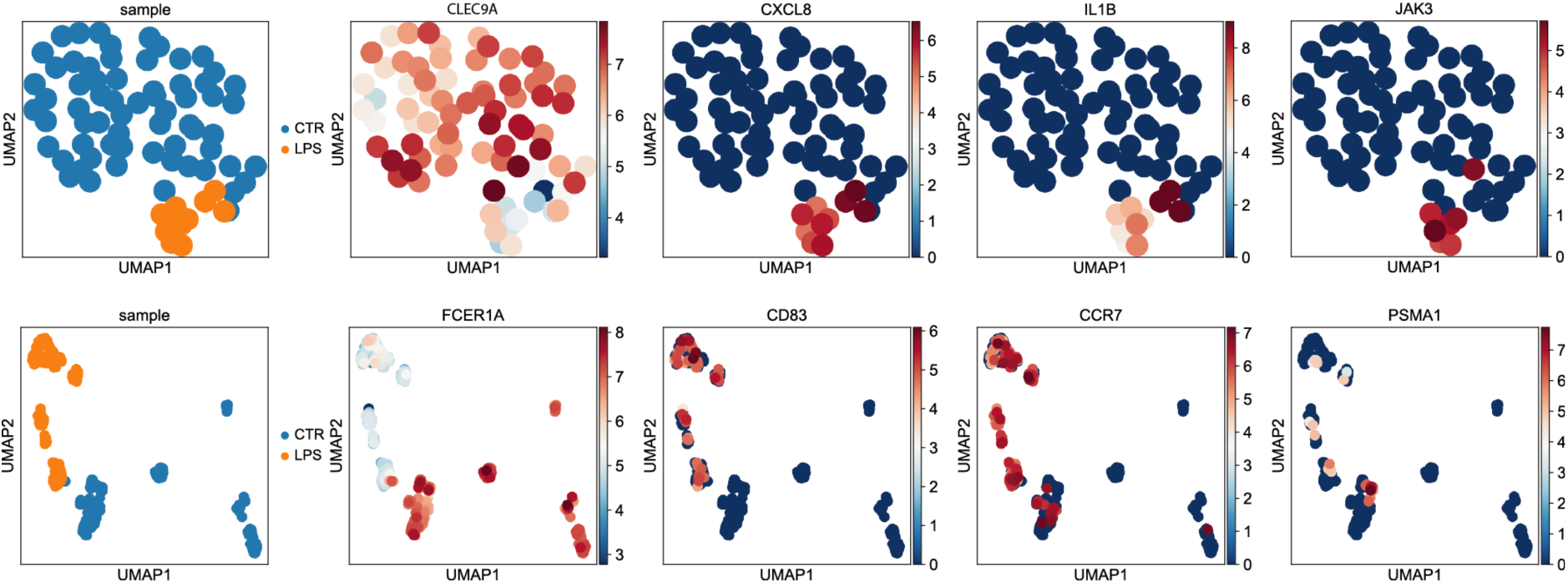
Two dendritic cell populations uniquely identified by C-DS. Top: UMAP plot showing DC1 cells, control (blue) and LPS (orange) stimulated, (Leiden r=0.5, n_neighbours = 20). While *CLEC9A+* expression is uniform in control and LPS stimulated cells, *CXCL8* and *IL1B* and *JAK3* expression can be detected exclusively in LPS stimulated cells. Bottom: UMAP plot showing DC2 cells, control (blue) and LPS (orange) stimulated. While subset marker, *FCER1A+* expression is high in control cells, the expression of *CD83*, *CCR7* and *PSMA1*, encoding DC activation is upregulated by LPS. Colour denotes gene expression level, as indicated by the heat-map legend (normalised UMI counts).

**Figure S7.**
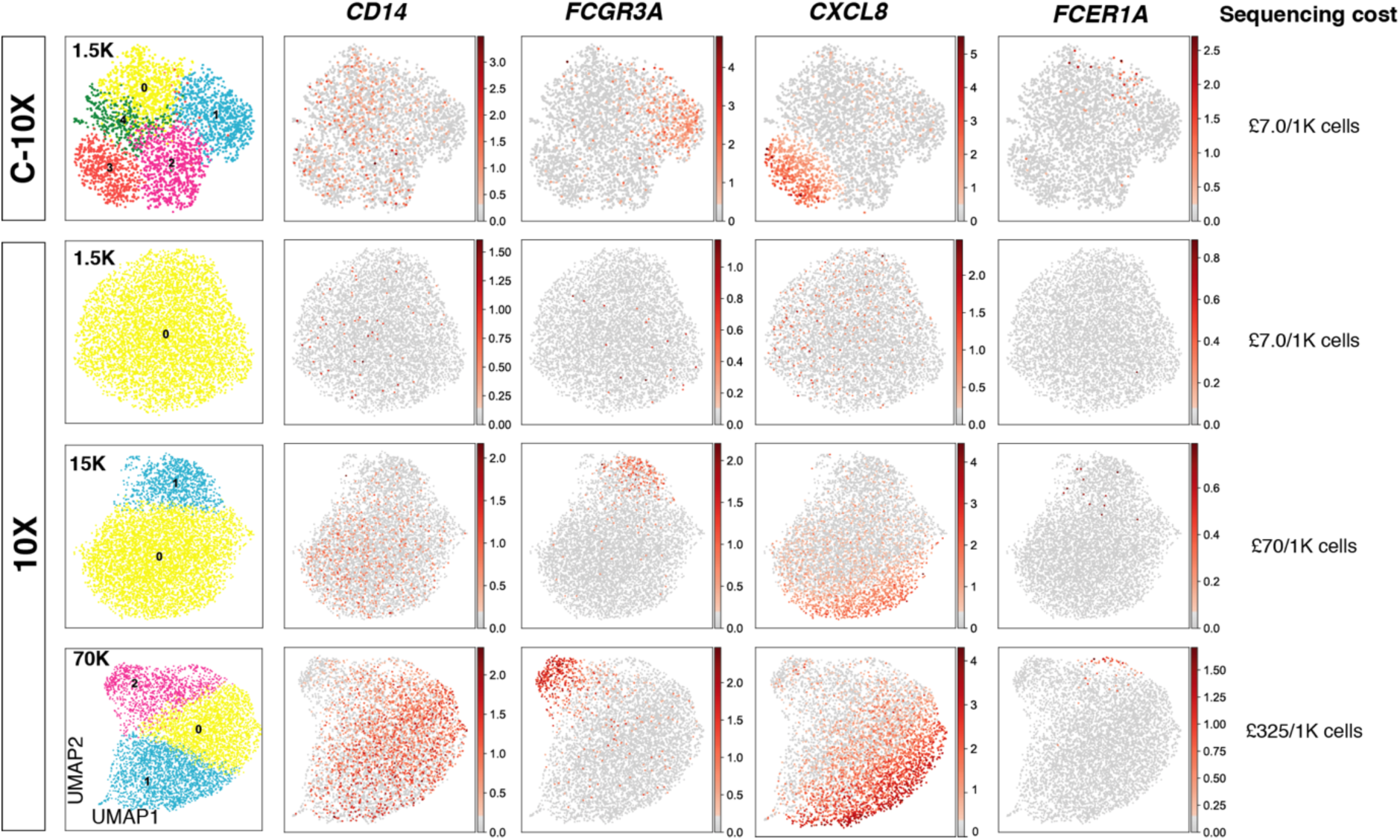
The Constellation method can be translated to other single-cell protocols. UMAP plots showing comparison of single cell sequencing of 6000 monocytes using C-10X at 1500 reads per cell sequencing depth with standard 10X at varying sequencing depths. Column 1: clustering results, Leiden r=0.5, n_neighbours = 20, columns 2-5: examples of monocyte activation expression markers. Colour denotes gene expression level, as indicated by the legend (normalised UMI counts). Right: the effective cost of sequencing 1000 cells.

**Figure S8.**
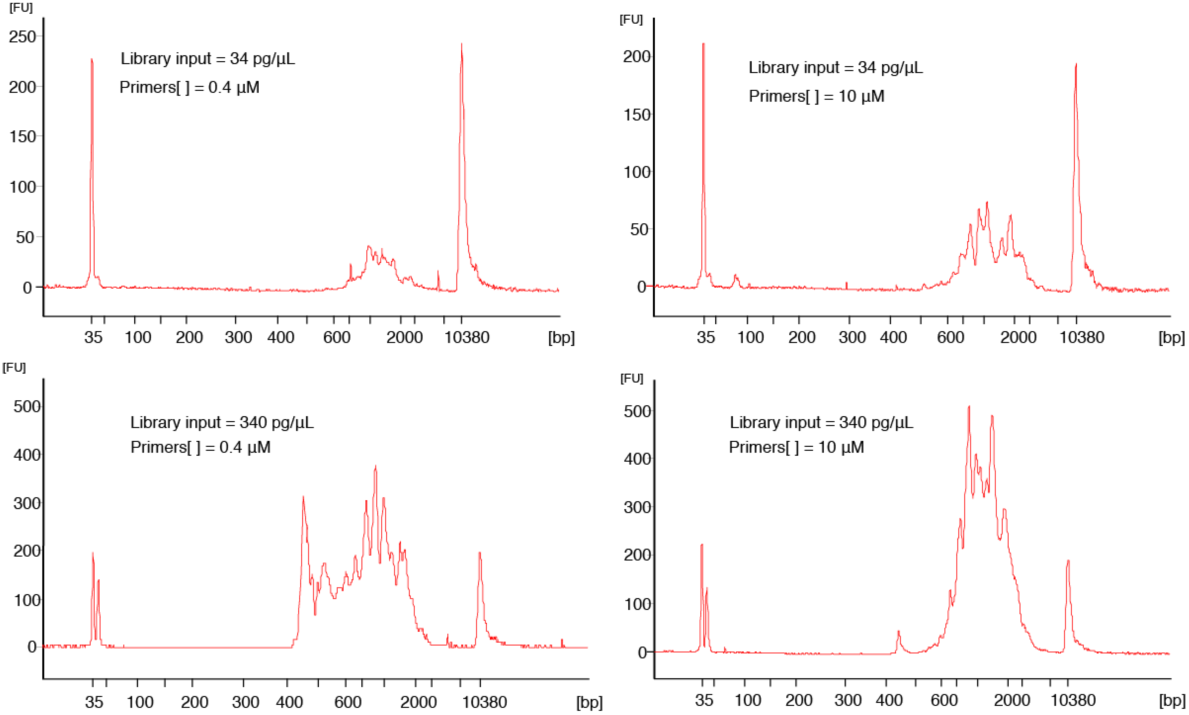
Optimization of Library preparation in C-10X. Typical plot from the bioanalyser (Agilent) showing the library input and primer concentration effect on library preparation for C-10X. Top: library input – 34 pg/mL, bottom – library input 340 pg/mL. Left: Primer concentration c= 0.4 ***μ***Mol, right: Primer concentration c= 10 ***μ***Mol. Y axis shows fluorescence units (FU) indicating signal intensity and product concentration. The spikes in the plot are characteristic for Constellation method due to the selection of targets with distinct molecular weights.

