## Supplementary material for "Resolving cellular systems by ultra-sensitive and economical single-cell transcriptome filtering": Table S4

### Supplementary Table 4

| | DropSeq | C-DropSeq | 10x genomics | C-10X |
|---|---|---|---|---|
| Encapsulation cost* | £5.50 | £5.50 | £135 | £135 |
| Library preparation* | £50 | £50 | £20 | £20 |
| Targeted amplification* | NA | £50 | NA | £20 |
| Sequencing* | £ 270** | £ 27*** | £ 270** | £ 27*** |
| **Total*** | **£325.50** | **£132.50** | **£425** | **£202** |
| Estimated labor time | 8 hours | 12 hours | 4 hours | 5 hours |

* Cost per 1000 cells
**at 50K reads/cell
***at 5K reads/cell
